# Active and predictive adjustment of pupil size for accommodating visual regularity

**DOI:** 10.1101/2025.04.01.646516

**Authors:** Jianrong Jia, Zhiming Kong, Fang Fang

## Abstract

The visual system efficiently encodes information by summarizing visual regularities that could be quantified by summary statistics. While summary statistics are thought to be processed in several cortical and subcortical areas, we investigated whether the earliest stage of visual processing, the pupil, is involved in this process. In three experiments, participants either performed ensemble orientation estimation with or passively viewed bar arrays with different bar orientation distributions while their pupil size was recorded. We found that pupil size increased when the orientation distribution became more dispersed and was closely linked to participants’ estimation performance. Furthermore, pupillary responses occurred automatically during passive viewing and could even predict ensemble orientation estimation performance. Moreover, when anticipating a dispersed distribution, the pupil dilated in advance. Our findings reveal a new cognitive role of the pupil - it actively and predictively adjusts its size to facilitate the extraction of visual regularity.

## Introduction

Information in the visual environment is structured in nature and contains a variety of regularities. These information regularities contribute to the efficiency of information coding, thus enabling pattern recognition and learning (Barlow, 2001; Kersten, 1987). Conversely, information that lacks regularity is hard to be used by the brain to form meaningful representations (Attneave, 1954). For example, to understand a complex scene consisting of multiple similar components, the visual system needs to statistically summarize incoming information from these components to extract regularities and form an ensemble perception (Alvarez, 2011). The mean value and variance of the component features in an ensemble are typically used to represent the regularity of the ensemble (Alvarez, 2011; Jia et al., 2022; Wang et al., 2023; Whitney and Yamanashi Leib, 2018). When the distribution of features is concentrated, the mean value is highly representative, making it easy for the visual system to extract the regularity. Conversely, when the distribution is dispersed, the mean value is less representative and acquiring the regularity is challenging (Semizer and Boduroglu, 2021).

It has been suggested that visual neurons need to pool together information over (at least part of) the visual field to form the ensemble representation through feedforward, feedback, and even horizontal connection mechanisms (Epstein et al., 2020; Parkes et al., 2001; Utochkin et al., 2023). Ensemble statistics (e.g., mean, variance) have been demonstrated to be represented in multiple brain regions along the visual information processing hierarchy. Neuroimaging studies found that ensemble orientation was represented in early visual cortical areas, V2 and V3, as well as the intraparietal sulcus (IPS) and superior parietal lobule (SPL) (Tark et al., 2021). Object ensemble was represented in the parahippocampal place area (PPA) (Cant et al., 2015; Cant and Xu, 2012), and face ensemble was represented in the IPS and superior frontal gyrus (SFG) (Im et al., 2017). Surprisingly, recent psychophysical studies using the eye-of-origin paradigm implied that ensemble size could be even represented as early as in subcortical brain structures (Zeng et al., 2024; Zhao et al., 2023), though neuronal receptive fields in these structures are quite small.

The pupil, the earliest part of the visual pathway, which is traditionally believed to respond to simple luminance changes by adjusting its size, has recently been shown to engage in several high-level cognitive processes (Grujic et al., 2024; Joshi and Gold, 2020). Pupil size is considered as an indirect, non-invasive measure of norepinephrine (NE) release (Aston-Jones and Cohen, 2005; Joshi et al., 2016). Norepinephrine is generated in the brainstem nucleus locus coeruleus (LC) and projects to multiple brain areas (Joshi and Gold, 2020; Strauch et al., 2022). The LC is thought to play a critical role in representing the statistical structure of environmental information (Zhao et al., 2019a) and updating the internal models of the brain (Zénon, 2019). Therefore, it is intriguing to ask whether and how the brain’s processing of the statistical regularities of visual information is reflected in pupil size.

In this study, we presented human participants with bar arrays matched for physical luminance and position and manipulated the orientation of the bars such that the bar arrays exhibited different statistical distributions (i.e., difference variances or regularities) in orientation. Participants either performed statistical summarization on the bar arrays (i.e., ensemble orientation estimation) or passively viewed them while their pupil size was recorded. We observed that the ensemble orientation estimation was worse when the bar orientation distribution was dispersed, compared to when it was concentrated. Critically, pupil size was significantly larger in the dispersed than in the concentrated distribution and was associated with the ensemble orientation perception. This pupillary response occurred automatically during passive viewing of the bar arrays and could predict individual participants’ ensemble orientation estimation performance. Furthermore, when a bar array with a dispersed distribution was anticipated, the pupil became larger before the array presentation than when an array with a concentrated distribution was anticipated. These findings highlight the active and predictive function of pupillary responses in acquiring visual information, summarizing visual statistics, and accommodating visual regularity.

## Results

### Bar orientation distribution affects ensemble perception and pupil size

In Experiment 1, we presented participants with 12 bars in the peripheral visual field (Fig. 1A). Participants were instructed to estimate the mean orientation of the bar array (i.e., ensemble orientation) with the method of adjustment (Fig. 1B). We manipulated the distribution of the orientations of the 12 bars such that four orientation distributions with different degrees of centralization were formed. Specifically, individual bars in an array were assigned with one of four orientations, i.e., 10°, 30°, 50°, and 70° clockwise/counter-clockwise relative to the vertical orientation, forming four orientation distributions — ‘9111’, ‘6321’, ‘4332’, and ‘3333’, with each digit representing the number of bars in one orientation in the array. For instance, in the distribution ‘9111’, nine of the 12 bars were presented in one orientation, and the other three bars were presented in each of the other three orientations. Notably, during the presentation, all the 12 bars were jittered in ±2° of their assigned orientations so that no bars were exactly the same. Participants’ pupillary responses were recorded during the presentation of the array.

**Fig. 1.**
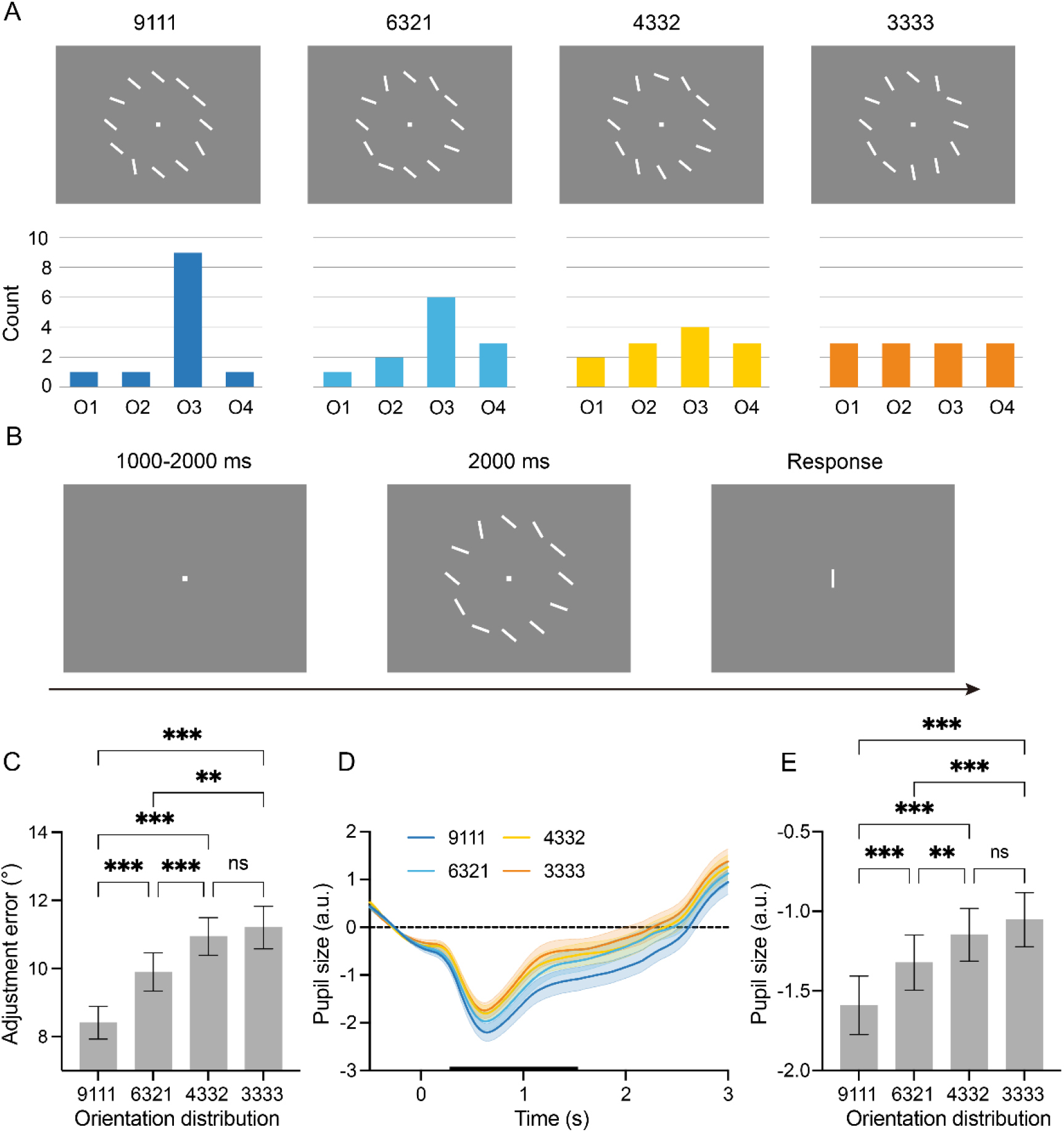
Procedure and results of Experiment 1. **A** (Top) Demonstrations of four orientation distributions used in Experiment 1. The bars were presented in four orientations. In the distribution ‘9111’, nine of the 12 bars were presented in one orientation, and the other three bars were presented in each of the other three orientations. The 12 bars in distributions ‘6321’, ‘4332’, and ‘3333’ were assigned to the four orientations in a ratio of 6:3:2:1, 4:3:3:1, and 3:3:3:3, respectively. (Bottom) Histograms show the number of bars in the four orientations. O1-O4 indicate the four orientations. **B** Experimental procedure of a full trial. Each trial began with the presentation of a fixation point followed by a bar array. The array consisted of 12 white bars displayed on a gray background. A single probe bar was presented subsequently. Participants adjusted the orientation of the probe bar to reproduce the mean orientation of the 12 bars in the array. **C** Adjustment errors are expressed as the difference between the reproduced orientation and the mean orientation of the bar array for the four orientation distributions. **D** Pupillary responses to bar arrays for the four orientation distributions. The black bar on the horizontal axis indicates the time period used for the statistical test. **E** The mean pupil sizes during the pupil constriction period (0.3-1.5 s) for the four orientation distributions. The shaded areas and error bars denote the standard error of the mean (SEM) across participants. *** indicate P < 0.001, ** indicate P < 0.01, ns indicate P > 0.05. All paired comparisons were Bonferroni corrected.

We first investigated whether the bar orientation distribution affected participants’ ensemble orientation estimation. Figure 1C showed that there was a significant difference in the adjustment error between the four orientation distributions (F(3,78) = 33.01, P < 0.001, partial η^2^ = 0.56). Interestingly, with the orientation distribution changing from centralized (i.e., ‘9111’) to uniform (i.e., ‘3333’), the adjustment error increased monotonically. Specifically, the adjustment error for the distribution ‘9111’ was significantly smaller than those for the other three distributions (all ts > 4.76, all Ps < 0.001, and all Cohen’s ds > 0.51). The adjustment error for the distribution ‘6321’ was significantly smaller than those for distributions ‘4332’ and ‘3333’ (both ts > 3.34, both Ps < 0.008, and both Cohen’s ds > 0.36). There was no significant difference in adjustment error between distributions ‘4332’ and ‘3333’ (t(26) = 0.83, P = 1, Cohen’s d = 0.09).

Pupillary responses to the four orientation distributions are shown in Fig. 1D. After stimulus onset, the pupil constricted rapidly, exhibiting a typical phasic response to the presentation of the array. Importantly, while the physical luminance was matched across the four different orientation distributions, pupil constriction was the largest in the most centralized distribution (i.e., ‘9111’) and the smallest in the most uniform distribution (i.e., ‘3333’). For the purpose of statistical analyses, we calculated the mean pupil sizes during the pupil constriction period (0.3-1.5 s after stimulus onset). We found a monotonic relationship between the mean pupil size and the centralization degree of the distribution (Fig. 1E). First, there was a significant difference in pupil size between the four orientation distributions (F(3,78) = 50.66, P < 0.001, partial η^2^ = 0.66). Furthermore, the pupil constriction for the distribution ‘9111’ was significantly greater than those for the other three distributions (all ts > 5.72, all Ps < 0.001, and all Cohen’s ds > 0.29), and the pupil constriction was significantly greater for the distribution ‘6321’ than those for distributions ‘4332’ and ‘3333’ (both ts > 3.72, both Ps < 0.002 and both Cohen’s ds > 0.19). There was no significant difference in pupil constriction between distributions ‘4332’ and ‘3333’ (t(26) = 2.01, P = 0.289, Cohen’s d = 0.11). These results mirrored the ensemble orientation estimation results.

To further quantify the relationship between the bar orientation distribution, the ensemble orientation perception, and the pupil size, we used entropy from the information theory to quantify the information amount of the orientation distribution, with smaller (larger) entropy indicating a more concentrated (dispersed) distribution and stronger (weaker) representativeness of the mean value in the distribution. We examined the relationship between the entropy of the orientation distribution and the behavioral performance and the pupil size using two linear mixed models, in which the entropy of the orientation distribution was used as the fixed-effect factor and participants as the random-effect factor to predict the behavioral performance and pupil size. Figure 2A showed that the entropy of the orientation distribution could predict the adjustment error (β = 3.46, P < 0.001), with the greater the entropy (the more dispersed the orientation distribution) the larger the adjustment error (Fig. 2A). Furthermore, the entropy of the orientation distribution could also predict the pupil size. Significant prediction (P < 0.05, FDR corrected) was observed starting from 0.25 s after stimulus onset (Fig. 2B), with greater entropy predicting smaller pupil constriction (i.e., larger pupil size).

**Fig. 2.**
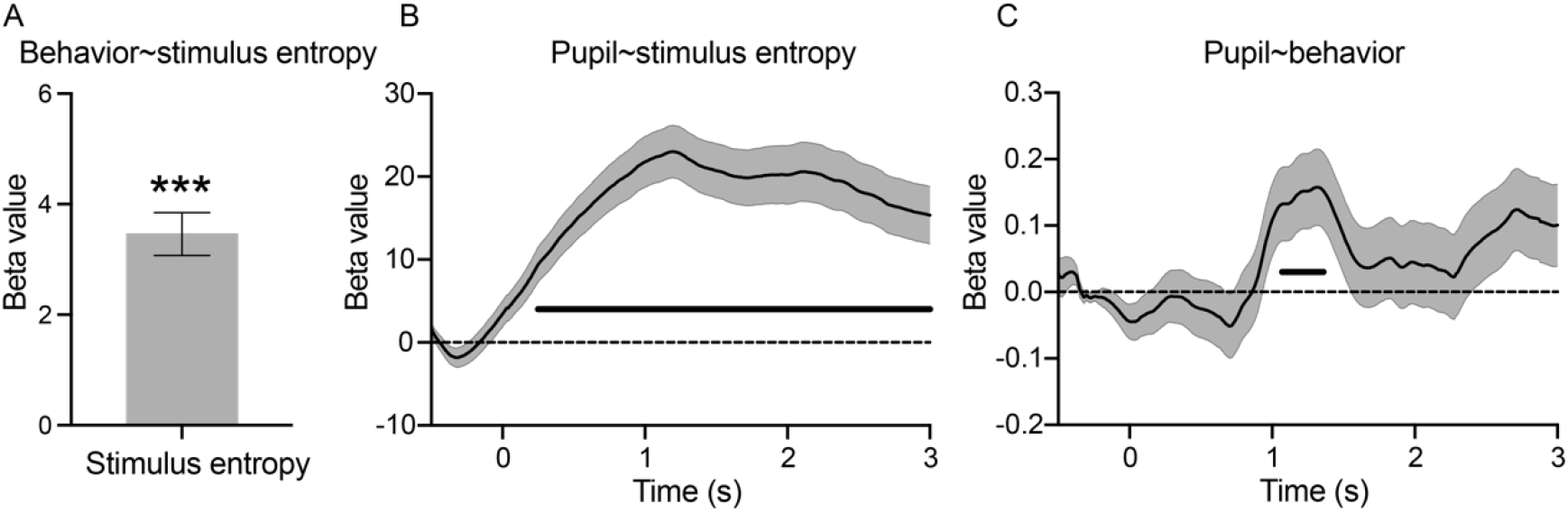
Relationship between entropy of orientation distribution, ensemble perception performance, and pupil size in Experiment 1. **A** Beta value of entropy in the prediction of adjustment error. **B** Beta values of entropy in the prediction of pupil size at each time point. **C** Beta values of adjustment error in the prediction of pupil size at each time point. The horizontal thick bars in **B** and **C** show the significant time points (P < 0.05) after FDR correction. The shaded areas and error bars denote the SEM across participants. *** indicate P < 0.001.

In the third linear mixed model, we used the adjustment error as the fixed-effect factor and participants as the random-effect factor to predict the pupil size at each time point. Figure 2C showed that the adjustment error could predict (P < 0.05, FDR corrected) the pupil size at 1.07-1.36 s after stimulus onset (Fig. 2C). Larger behavioral adjustment error predicted smaller pupil constriction (i.e., larger pupil size). These results showed the close relationship between the bar orientation distribution and the ensemble perception and the pupil size, and further demonstrated that the entropy of the information in the bar array was reflected in participants’ ensemble perception and pupillary responses.

### Pupil size reflects current bar orientation distribution rather than distribution switching

Pupil dilation was found to occur when stimuli switched from regular to irregular distribution instead of the other way around (2019a), which might reflect violation of regularity instead of perception of irregularity as suggested above. To examine this issue, we categorized the four distributions into two groups based on their degree of regularity (high regularity: ‘9111’ and ‘6321’; low regularity: ‘4332’ and ‘3333’), and examined the effect of switching between different degrees of regularity on pupillary responses. Fig. 3A shows pupillary responses for high or low regularity trials, which were immediately preceded by a high or low regularity trial. For instance, ‘L-H’ indicates that the current trial has high regularity, while the preceding trial has low regularity.

**Fig. 3.**
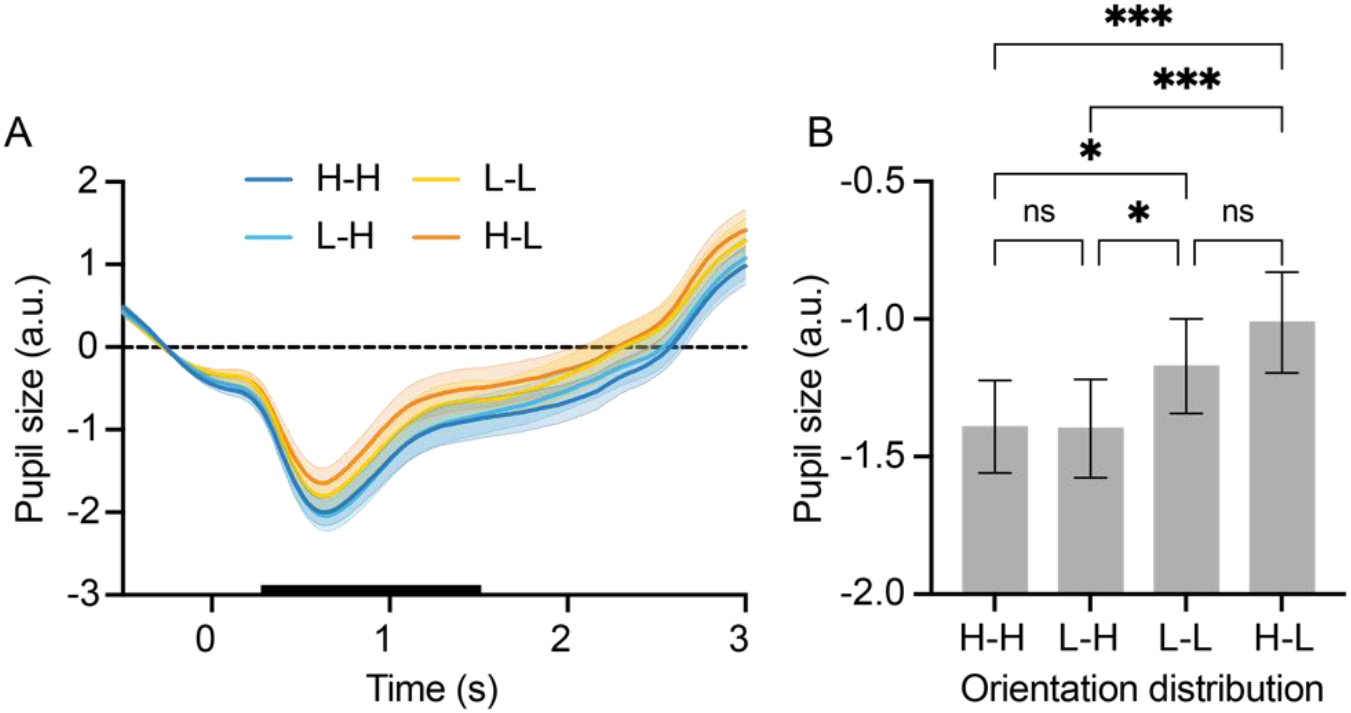
Pupillary responses to the switching of orientation distribution. **A** Pupillary responses to the four types of orientation distribution switching. ‘H-H’ represents the switching from high regularity (preceding trial) to high regularity (current trial), ‘L-H’ the switching from low regularity to high regularity, ‘L-L’ the switching from low regularity to low regularity, and ‘H-L’ the switching from high regularity to low regularity. The black bar on the horizontal axis represents the time period used for the statistical test (B). **B** The mean pupil sizes during the pupil constriction period (0.3-1.5 s) for the four conditions. The shaded areas and error bars denote the SEM across participants. *** indicate P < 0.001, * indicate P < 0.05, ns indicate P > 0.05. All paired comparisons were Bonferroni corrected.

For statistical analysis (Fig. 3B), we extracted the mean pupil sizes during the pupil constriction period (0.3-1.5 s after stimulus onset). The 2 (regularity of the current trial) * 2 (regularity of the preceding trial) repeated-measure ANOVA showed a significant main effect of the regularity of the current trial (F(1,26) = 26.88, P < 0.001, partial η^2^ = 0.51), but not the regularity of the preceding trial (F(1,26) = 2.63, P = 0.117, partial η^2^ = 0.09). The interaction effect was not significant (F(1,26) = 2.56, P = 0.122, partial η^2^ = 0.09). Notably, we replicated Zhao et al.’s finding that the pupil size was larger in the ‘H-L’ condition than in the ‘L-H’ condition (t(26) = 5.19, P < 0.001, Cohen’s d = 0.42). However, only this finding cannot prove that the observed pupil size changes merely reflect statistical distribution switching. Instead, when comparing switched with unswitched conditions, we found no significant difference in pupil size as long as the regularity of the current array was matched (‘L-H’ vs. ‘H-H’: t(26) = 0.09, P = 1.000, Cohen’s d = 0.01; ‘L-L’ vs. ‘H-L’: t(26) = 2.14, P = 0.214, Cohen’s d = 0.17). Therefore, the changes in pupil size reflect the perception of the current distribution rather than the distribution switching.

### Pupillary response is not due to attentional state

One possible explanation for the greater pupil constriction to the distribution ‘9111’ is the existence of singular orientations (i.e., the three ‘1’s) in the distribution that might have induced changes in the attentional state due to their high visual salience. We used participants’ microsaccade inhibition as a measure of attentional state to test this possibility. Stimulus presentation usually causes the inhibition of microsaccade, as evidenced by a reduction of microsaccade rate, and the magnitude of the reduction was found to be modulated by attention (Contadini-Wright et al., 2023; Valsecchi et al., 2007; Zhao et al., 2019b). Here, presentation of all four orientation distributions indeed elicited significant microsaccade inhibition (Fig. 4A). However, the averaged microsaccade rate over the inhibition time period (0.05-0.14 s after stimulus onset) (Fig. 4B) did not show any significant difference among the four distributions (F(3,78) = 1.56, P = 0.205, partial η^2^ = 0.06), suggesting that the four orientation distributions did not elicit different attentional state changes. This result is consistent with a recent finding that pupil size does not respond to stimulus salience (Zhao et al., 2019b).

**Fig. 4.**
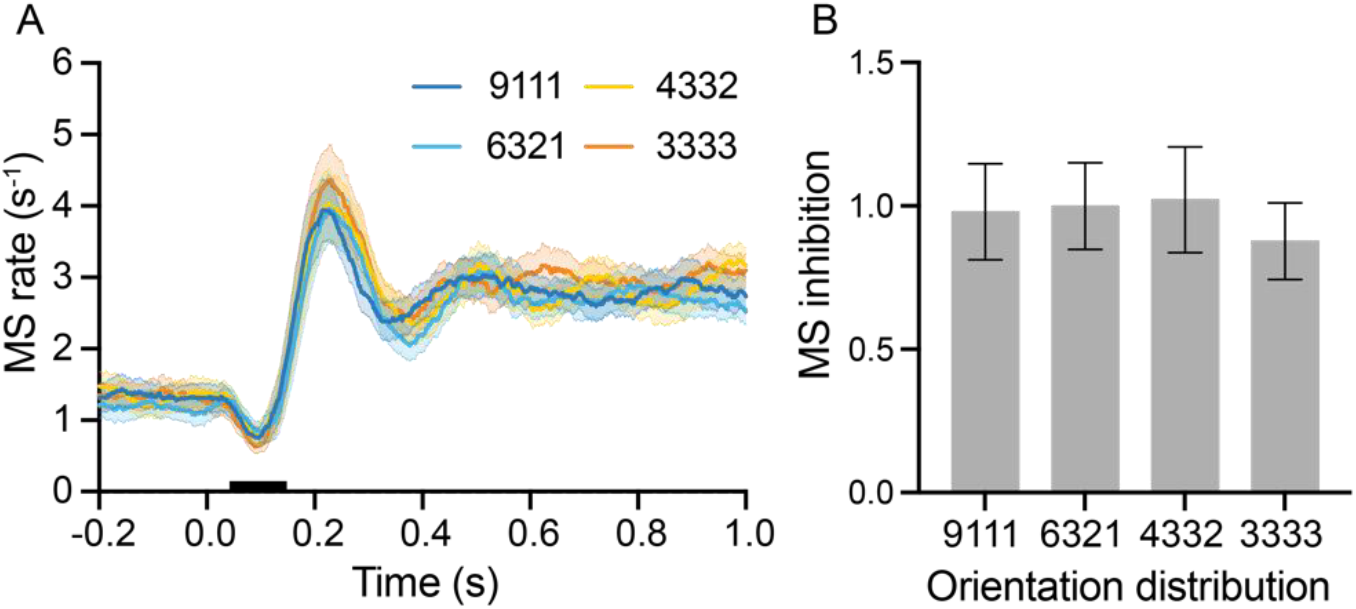
Microsaccade responses for the four orientation distributions. **A** Microsaccade rates during the presentation of the bar array for the four orientation distributions. The black bar on the horizontal axis represents the time period used for the statistical test (B). **B** Microsaccade inhibition was expressed as the mean microsaccade rate during the inhibition time period (0.05-0.14 s). The shaded areas and error bars denote the SEM across participants.

### Close association between behavioral adjustment error during task and pupil size during passive viewing

Another explanation for the pupillary responses in Experiment 1 is that the pupil size could be merely modulated by the effort participants took in the ensemble orientation estimation task. To test this possibility, we investigated whether the pupil size changes to the different orientation distributions could occur automatically without the requirement for performing the task and whether the automatic pupillary response could even reflect individual differences in the task performance. To achieve this goal, Experiment 2 consisted of an ensemble orientation estimation task phase which was similar to that in Experiment 1 and a passive viewing phase where participants were exposed to bar arrays with different orientation distributions without any task. The bar arrays were the same as those in Experiment 1.

Figure 5A showed that there were significant differences in the adjustment error of the ensemble orientation estimation task between the four orientation distributions (F(3,81) = 20.23, P < 0.001, partial η^2^ = 0.43), replicating the results in Experiment 1. Specifically, the adjustment error for the distribution ‘9111’ was significantly smaller than those for the other three distributions (all ts > 3.86, all Ps ≤ 0.001, and all Cohen’s ds > 0.34). The adjustment error for the distribution ‘6321’ was significantly smaller than that of the distribution ‘3333’ (t(27) = 3.43, P = 0.006, Cohen’s d = 0.31) but not that of the distribution ‘4332’ (t(27) = 2.09, P = 0.240, Cohen’s d = 0.19). There was no significant difference in adjustment error between distributions ‘4332’ and ‘3333’ (t(27) = 1.34, P = 1, Cohen’s d = 0.12).

**Fig. 5.**
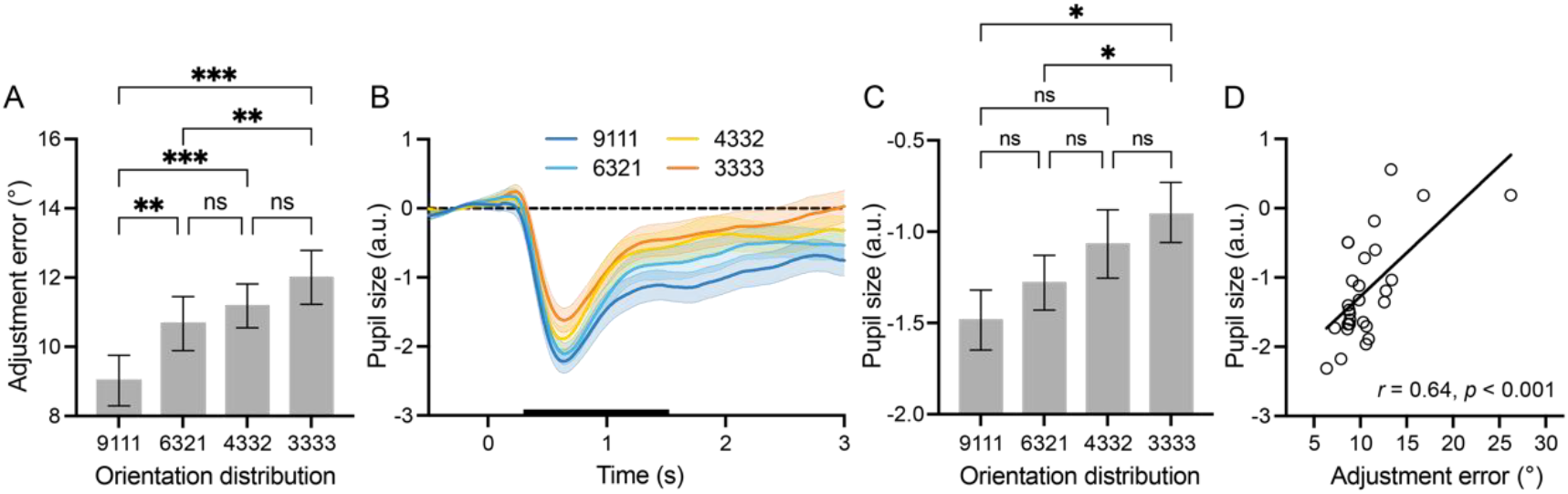
Behavioral and pupillary results of Experiment 2. **A** Adjustment errors for the four orientation distributions. **B** Pupillary responses to bar arrays for the four orientation distributions. The black bar on the horizontal axis represents the time period used for the statistical test. **C** The mean pupil sizes during the pupil constriction period (0.3-1.5 s) for the four orientation distributions. **D** Pearson correlation between the adjustment error of the ensemble orientation estimation task and the pupil size during passive viewing across participants. The shaded areas and error bars denote the SEM across participants. *** indicate P < 0.001, ** indicate P < 0.01, * indicate P < 0.05, ns indicate P > 0.05. All paired comparisons were Bonferroni corrected.

Pupillary responses to the four orientation distributions during passive viewing were shown in Fig. 5B. Replicating Experiment 1, pupil constriction was the largest in the most centralized distribution (i.e., ‘9111’) and the smallest in the most uniform distribution (i.e., ‘3333’). We calculated the mean pupil sizes during the pupil constriction period (0.3-1.5 s after stimulus onset). As shown in Fig. 5C, there were significant differences in pupil size between the four orientation distributions (F(3,81) = 5.69, P = 0.001, partial η^2^ = 0.17). Furthermore, the pupil constriction for the distribution ‘9111’ was significantly greater than that for the distribution ‘3333’ (t(27) = 3.34, P = 0.015, Cohen’s d = 0.66) but not those for distributions ‘6321’ and ‘4332’ (both ts < 2.82, both Ps > 0.054, and both Cohen’s ds < 0.48), and the pupil constriction was significantly greater for the distribution ‘6321’ than that for the distribution ‘3333’ (t(27) = 2.93, P = 0.041, Cohen’s d = 0.43) but not for the distribution ‘4332’ (t(27) = 1.35, P = 1.000, Cohen’s d = 0.24). There was no significant difference in pupil constriction between distributions ‘4332’ and ‘3333’ (t(27) = 1.36, P = 1, Cohen’s d = 0.19). Next, for each participant, we averaged their adjustment errors and pupil sizes across the four orientation distributions. Then we calculated the Pearson correlation between the averaged adjustment error and the averaged pupil size across participants and found that they were significantly and positively correlated (r = 0.64, P < 0.001) (Fig. 5D). These results suggest, on the one hand, that the pupillary responses to different orientation distributions are automatic, while on the other hand, participants who had lower sensitivity in estimating the ensemble orientation had less automatic pupil constriction during passive viewing. The automatic pupillary responses thus reflect the statistical summarization ability at the individual participant level.

### Pupil size reflects anticipated bar orientation distribution

In Experiments 1 and 2, we found that for dispersed bar orientation distributions, the pupil automatically increased its size, and that the pupil size was associated with the ensemble perception within and across participants. A possible explanation is that, for dispersed orientation distributions, the visual system actively dilates the pupil size and gathers more information about individual orientations (Mathôt, 2018; Vilotijević and Mathôt, 2023) in order to statistically summarize them.

In Experiment 3, we explored whether the pupil could respond predictively when anticipating the appearance of bar arrays with different distributions. We presented participants with a horizontal or vertical bar as a cue (Fig. 6A), followed by a bar array. In the vertical cue trials, 80% of the bar arrays were the ‘9111’ bar array and the remaining 20% were the ‘3333’ bar array. In contrast, in the horizontal cue trials, 80% of the bar arrays were the ‘3333’ bar array and the remaining 20% were the ‘9111’ bar array. This procedure rendered participants to generate anticipation of one of the two types of bar arrays according to the cue. The ‘9111’ and ‘3333’ bar arrays were identical to those in Experiment 1 and participants were instructed to passively view them. We investigated whether the pupil could predictively adjust its size to prepare for stimulus processing when participants anticipate bar arrays with different orientation distributions.

**Fig. 6.**
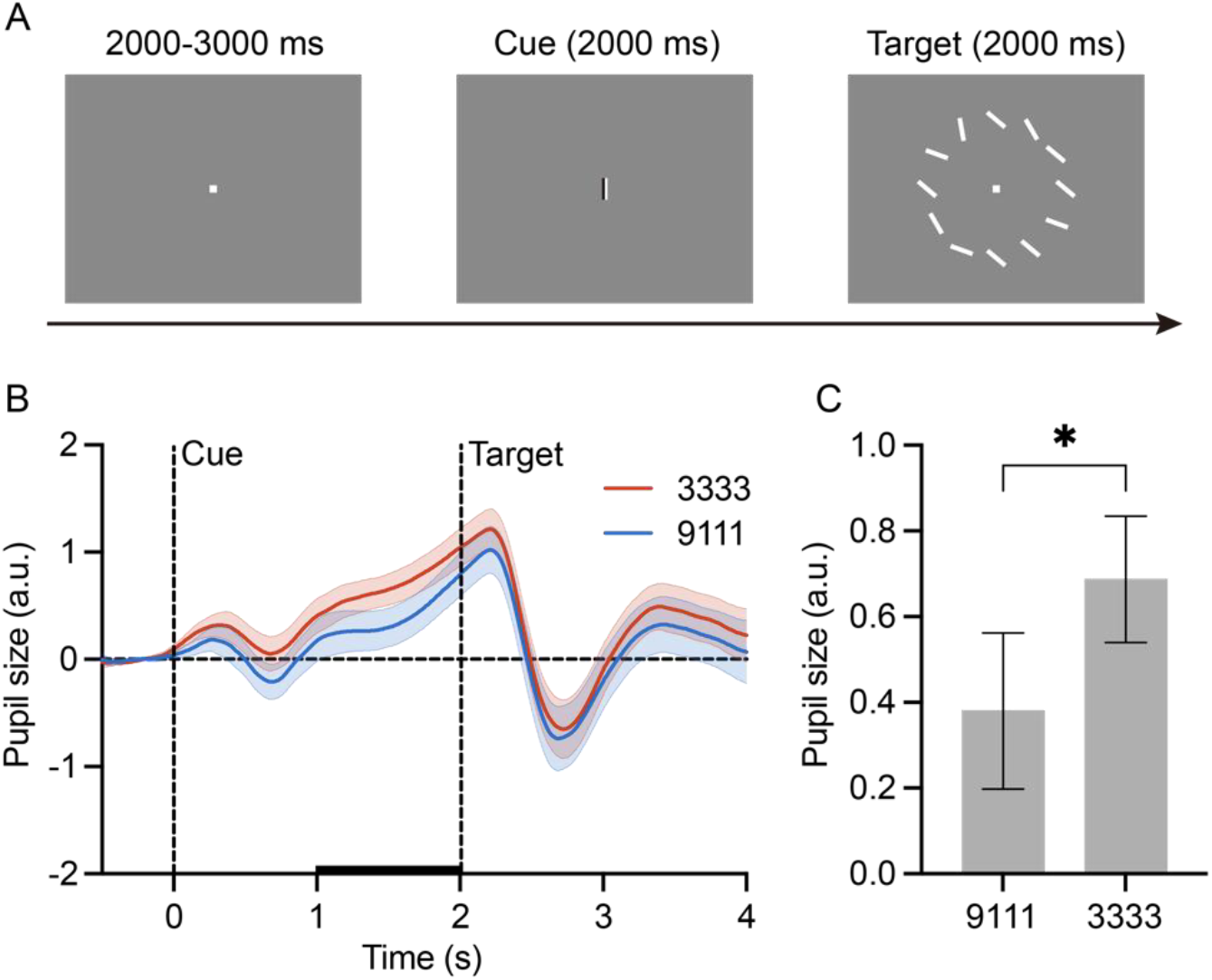
Procedure and pupillary responses of Experiment 3. **A** Experimental procedure. A half-black and half-white vertical or horizontal bar was presented as a cue for 2 s. In the vertical cue trials, the ‘9111’ bar array was presented in 80% trials and the ‘3333’ bar array was presented in the remaining 20% trials. In contrast, in the horizontal cue trials, the ‘3333’ bar array was presented in 80% trials and the ‘9111’ bar array was presented in the remaining 20% trials. The bar array (i.e., target) was presented for 2 s, followed by the next trial after a random interval between 2-3 s. Participants maintained their fixation and passively viewed the bar array. **B** Pupillary responses to the vertical and horizontal cues. The black bar on the horizontal axis represents the time period used for the statistical test. **C** Mean pupil sizes of 1 s before the target onset. The shaded areas and error bars denote the SEM across participants. * indicate P < 0.05.

Figure 6B shows that the pupil size was smaller when the ‘9111’ bar array was anticipated than when the ‘3333’ bar array was anticipated. We averaged the pupil size 1 s before the bar array onset for statistical comparison (Fig. 6C). The result confirmed our impression (t(28) = 2.42, P = 0.022, Cohen’s d = 0.45), demonstrating that the pupil can make adjustment to prepare for processing anticipated stimuli.

## Discussion

This study investigated the role of the pupil in processing summary statistics and regularities of visual stimuli. We demonstrated that pupil size was modulated by the statistical distribution of visual stimuli and was related to the perceptual summary (e.g., ensemble orientation estimation) of these stimuli. Specifically, the pupil size was smaller when the visual stimuli had a concentrated distribution and was larger when the stimuli had a dispersed distribution. These pupillary responses were automatic, independent of the task and attentional state. Linear mixed models and Pearson correlation analyses revealed that the pupillary responses reflected individual sensitivity in the perceptual summary. Furthermore, the pupil responded predictively to the anticipated statistical distribution before stimulus presentation. Taken together, these findings demonstrated that the brain’s processing of summary statistics and regularities of visual stimuli is reflected in adjusting pupil size in an active and predictive manner.

We believe that this study provides the first conclusive evidence for the pupil’s active role in summarizing visual information and accommodating visual regularity. Previous researches have shown that pupil size could be modulated by various statistical events and structures, including stimulus probability (Qiyuan et al., 1985), violation of learned statistical regularity (Zhao et al., 2019a), timing of stimulus appearance (Schwiedrzik and Sudmann, 2020), precision of auditory statistical distribution (Silvestrin et al., 2021), decision-making uncertainty (Preuschoff, 2011; Urai et al., 2017), and the ambiguity (Graves et al., 2021) and predictability (Milne et al., 2021) of stimulus. Together with these findings, here we propose a new functional role of the pupil in statistical summarization of visual information. Specifically, when a stimulus is uncertain, ambiguous, unpredictable, or contains dispersedly distributed components, the pupil is informed by the perceived regularity of the stimulus and actively dilates to gather more information in the visual field. Therefore, the visual system could summarize the information and capture the regularity in a more precise way. Through this interacting process between pupil size adjustment and regularity extraction, the visual system may finally reach an optimal state for encoding the summary statistics and regularity.

Even more interestingly and surprisingly, our results suggest that the pupillary response could even reflect the prediction of visual regularity, aligning with predictive coding models of visual processing (Huang and Rao, 2011; Rao and Ballard, 1999). The brain learns statistical regularities in the environment and predicts upcoming stimuli. Understanding and predicting stimuli can largely modulate the strength and pattern of neural responses in the brain (Carlsson et al., 2000; Murray et al., 2004, 2002; Rao and Ballard, 1999). In complex and dynamic environments, the brain must not only adjust its own responses but also coordinate sensory organs based on knowledge and prediction in order to respond actively and predictively to stimuli. In our study, according to the cue information, the visual system transmitted the prediction of the bar orientation distribution to brain areas controlling the pupil size in a top-down manner (Huang and Rao, 2011; Rao and Ballard, 1999). The pupil’s predictive adjustment before stimulus presentation aligns with previous findings on the primary visual cortex engaging in preplaying anticipated visual events (Ekman et al., 2017; Peelen et al., 2023). Our work significantly extends this predictive view by demonstrating a similar function at the earliest stage of the visual processing stream (i.e., the pupil).

It could be argued that, because stimuli with different statistical distributions may elicit different magnitudes of salience that could affect pupil size consequently, the observed pupillary responses might be simply explained as an attentional effect (Strauch et al., 2022; Wang et al., 2014). However, we believe that this explanation is very unlikely. First, we observed pupil size changes even during passive viewing (without attentional and task demand) that could predict participants’ ensemble orientation estimation performance. Second, we used microsaccade inhibition, a direct measure of attentional state (Contadini-Wright et al., 2023; Valsecchi et al., 2007; Zhao et al., 2019b), and found no significant difference between statistical distributions.

Summary statistics of visual features have been suggested to be represented in cortical (Cant and Xu, 2012; Im et al., 2017; Tark et al., 2021) and even subcortical brain areas (Zeng et al., 2024; Zhao et al., 2023). Our study reveals that the pupil also participates in statistical summary processing. Pupil size was found to be mainly controlled by several subcortical structures, including locus coeruleus, cholinergic basal forebrain, and colliculi (Joshi et al., 2016; Lloyd et al., 2023), implying a common subcortical neural system involved in both statistical summary processing and pupil size control. It should be noted that top-down modulation from the cortex can influence subcortical pupil control circuits (Aston-Jones and Cohen, 2005). Future studies using neuronal recording techniques should be performed to dissect cortical and subcortical circuits through which the statistical summary representation affects pupil size.

Pupillary responses have been suggested to be linked to the activity of the LC-NE system (Joshi et al., 2016). This system has two modes, tonic and phasic, and their balance reflects the exploration/exploitation trade-off in cognitive processing (Aston-Jones and Cohen, 2005), which can be reflected in pupil size changes (Gilzenrat et al., 2010; Jepma and Nieuwenhuis, 2011). Specifically, the switching of cognitive processing from exploitation to exploration is accompanied by pupil dilation. To some extent, our findings can be explained by this mechanism. Participants showed greater pupil constriction, i.e., smaller pupil sizes, for bar arrays with concentrated orientation distributions. Conversely, dispersed distributions led to less pupil constriction. Because the knowledge contained in concentrated distributions is regular and clear, cognitive processing may favor exploitation over exploration. In contrast, when the distribution is dispersed, the knowledge is irregular and unclear, and cognitive processing may instead favor exploration.

In conclusion, by combining psychophysical and pupillary response measures, we demonstrate that the pupil could adjust its size in response to different summary statistics and regularities in an active and predictive way. Beyond passively reflecting input luminance changes as traditionally believed, this previously unknown or largely unexplored function of the pupil can undoubtedly play a key role in acquiring necessary visual information, summarizing visual statistics, and accommodating visual regularity at the earliest stage of the visual processing stream. Our work will also prompt future research to further scrutinize the significance of pupillary responses in other visual processes, even cognitive processes more broadly.

## Materials and Methods

### Participants

A total of 84 adults (57 males, mean age: 21.3 years; 27 for Experiment 1, 28 for Experiment 2, 29 for Experiment 3) with normal or corrected-to-normal vision participated in the study. The sample size for Experiment 1 was determined based on a power analysis of repeated-measure ANOVA (effect size = 0.25, α = 0.05, power = 0.85, groups = 1, measurements = 4) performed prior to the execution of the experiment. For Experiments 2 and 3, the sample size was approximately the same as that for Experiment 1. The procedure of the study was approved by the local ethics committee (#20200629) and was in accordance with the Declaration of Helsinki. All participants gave written informed consent prior to the study.

### Apparatus and tools

MATLAB (The MathWorks) and the Psychophysics Toolbox (Brainard, 1997) were used to present the visual stimuli and to record behavioral responses. The experiments were performed in a dimly-lit and sound-proof laboratory. The visual stimuli were presented on a CRT monitor (resolution: 1024 × 768; refresh rate: 85 Hz). Participants were comfortably seated at a viewing distance of about 70 cm.

### Eye-movement recording

Eye movements were recorded binocularly at 1000 Hz with an EyeLink 1000 eye tracker (SR Research, Ottawa). To maintain good tracking accuracy, participants’ heads were stabilized with a chin rest. The eye tracker was calibrated with a standard 9-point calibration procedure at the beginning of the experiment and after every 40 trials.

### Experiment 1

#### Stimuli

The stimulus consisted of 12 white bars (1.29° × 0.32°, 104 cd/m^2^) evenly distributed along a virtual circle (6° radius) surrounding the center of a gray (23 cd/m^2^) screen. The orientations of the bars were categorized into two groups: clockwise and counter-clockwise, each consisting of four orientations. In the clockwise category, the four orientations were 10°, 30°, 50°, and 70° clockwise from the vertical orientation. In the counter-clockwise category, the four orientations were 10°, 30°, 50°, and 70° counterclockwise from the vertical orientation. We created four orientation distributions from the 12 bar orientations (Fig. 1A), which were labeled as ‘9111’, ‘6321’, ‘4332’, and ‘3333’ respectively. The digits in each label represented the number of bars in a specific orientation. For instance, in the distribution ‘9111’, nine out of the 12 bars were presented in one orientation, and the other three bars were presented in each of the other three orientations. The 12 bars in distributions ‘6321’, ‘4332’, and ‘3333’ were assigned to the four orientations in a ratio of 6:3:2:1, 4:3:3:1, and 3:3:3:3, respectively. For each trial, all the 12 bars were jittered in ±2° of their assigned orientations so that no bars were exactly the same. And all orientations were either in the clockwise or counter-clockwise category.

#### Procedure

In each trial, the bar array was displayed in the peripheral visual field for 2 s. Then a single probe bar was presented at the center of the screen. Participants were instructed to press keys to adjust the orientation of the probe bar to match the average orientation of the 12 bars in the preceding array (Fig. 1B). No time constraint was imposed during this adjustment process. Participants confirmed their adjustment result by pressing the ‘space’ key. The next bar array appeared after a 1-2 s interval. Participants were asked to maintain their fixation at the center of the screen throughout the experiment.

The experiment consisted of 8 blocks of 40 trials, resulting in 320 trials in total. Within each block, trials with the four orientation distributions were presented in random order. The experiment lasted approximately 45 minutes. Before the formal experiment, each participant completed 20 practice trials.

### Experiment 2

#### Stimuli

Same as in Experiment 1.

#### Procedure

Experiment 2 consisted of two phases. The procedure of the ensemble orientation estimation task phase was similar to that of Experiment 1. A bar array was presented for 0.5 s, followed by a single probe bar presented at the center of the screen. Participants were instructed to adjust the orientation of the probe bar to match the average orientation of the 12 bars in the preceding array. The next bar array appeared after a 1-2 s interval. In the passive viewing phase, a bar array was presented for 3 s, followed by a 1-2 s interval. The order of the two phases was counterbalanced across participants. Both phases consisted of four blocks of 64 trials, resulting in a total of 512 trials. Trials with the four orientation distributions were presented in a random order within each block. The experiment lasted approximately 45 minutes. Before the task phase, each participant completed 20 practice trials. Participants’ pupil data were recorded during the passive viewing phase.

### Experiment 3

#### Stimuli

The ‘9111’ and ‘3333’ bar arrays in Experiment 1 were used as the target stimulus. The cue was a half-black (0 cd/m^2^) and half-white (104 cd/m^2^) vertical or horizontal bar (1.29° × 0.32°) presented at the center of the screen.

#### Procedure

Each trial started with a vertical or horizontal bar, serving as a cue, presented at the screen center for 2 s. In the vertical cue trials, the ‘9111’ bar array was presented in 80% trials and the ‘3333’ bar array was presented in the remaining 20% trials. In contrast, in the horizontal cue trials, the ‘3333’ bar array was presented in 80% trials and the ‘9111’ bar array was presented in the remaining 20% trials. The bar array (i.e., target) was presented for 2 s, followed by the next trial after a random interval between 2-3 s. Participants maintained their fixation and passively viewed the bar array. The experiment consisted of 4 blocks of 60 trials, resulting in a total of 240 trials. Trials of two cueing conditions were randomly presented within each block. Participants completed 10 practice trials before the experiment. The experiment lasted approximately 30 minutes. Participants’ pupil data were recorded during passive viewing.

### Quantifying stimulus variability with entropy

We used entropy from information theory as a metric to measure the information amount of each bar orientation distribution:

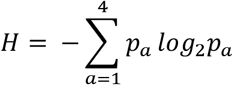

where H represents the entropy of an orientation distribution, *p*_*a*_ denotes the probability of a specific orientation in the distribution. Experiment 1 had four orientation distributions, so *a* ranges from 1 to 4. This classical measure of entropy can be conceptualized as a gauge of variability, uncertainty, or surprisal. A lower entropy value denotes that orientations in the distribution are less variable, while a higher entropy value denotes the opposite.

### Preprocessing of pupil data

The pupil data were preprocessed using the PuPl toolbox (Kinley and Levy, 2021), which had six steps. (1) Removal of outlier values: pupil data that were too large or too small, i.e., outside the range of Mean ± 2 SDs, and that dilated too rapidly were removed (Kret and Sjak-Shie, 2019). Additionally, data “islands” with a duration of less than 30 ms and an interval of more than 25 ms from adjacent data were excluded (Lemercier et al., 2014). (2) Removal of blinks: blinks were identified using the pupillometry noise method (Hershman et al., 2018). Since blinks appear as a sharp decrease and subsequent increase in measured pupil size, any data within 25 ms before and 100 ms after a detected blink were marked as blink data and removed as well. (3) Interpolation: missing pupil data were interpolated using a linear interpolation. The interpolation was constrained with the length of time to insert being within 500 ms and the difference of the pupil size between the two ends of the interpolated data being no more than one standard deviation. (4) Smoothing and downsampling: to eliminate high-frequency artifacts in the data, smoothing was performed with a 100-ms Hanning window, followed by downsampling to 100 Hz. (5) Defining epochs: data from 500 ms before the presentation of the bar array to 3000 ms after in Experiments 1 and 2 and data from 500 ms before the presentation of the cue to 4000 ms after in Experiment 3 were defined as an epoch. Any epoch containing more than 1% of missing pupil data was excluded. On average, 22 epochs were removed from each participant. Subsequently, data from the two eyes were averaged to create a single index. (6) Normalization and baseline correction: to mitigate the influence of individual differences in pupil size on the statistics of the data, the pupil data for each participant were converted to z-scores. Furthermore, post-stimulus data were baseline-corrected using the pre-stimulus 500 ms data as the baseline.

### Microsaccade detection

Microsaccades were identified using an improved version (Engbert and Mergenthaler, 2006) of the algorithm initially introduced by Engbert and Kliegl (2003). The process involved the mapping of horizontal and vertical eye positions into a velocity space, where microsaccades were detected by applying a velocity threshold of 5 standard deviations (Engbert and Mergenthaler, 2006). To be classified as a microsaccade, a saccade needed to satisfy certain criteria, including a temporal overlap between the two eyes, a minimum duration of 6 ms, and an amplitude below 1°. The microsaccade rate was examined in the time window from 0.2 s before the bar array onset to 1 s after using a rectangular moving window of 100 ms with steps of 1 ms (Engbert and Mergenthaler, 2006; Lv et al., 2022).

### Linear mixed models for the relationship between bar orientation distribution, behavioral performance, and pupil size

We employed three linear mixed models to assess the relationship between bar orientation distribution, behavioral performance, and pupil size. First, we fitted a linear mixed-effects model for the behavioral performance, with a fixed effect for the bar orientation distribution and a random effect for the participants. Next, we fitted a linear mixed-effects model for the pupil size at each time point, with a fixed effect for the bar orientation distribution and a random effect for the participants. Finally, we fitted a linear mixed-effects model for the pupil size at each time point, with a fixed effect for the behavioral performance and a random effect for the participants. The parameters of the linear mixed models were estimated with the maximum likelihood method, and the covariance matrix of the random effects was estimated with the Cholesky parameterization.

### Statistics

Comparisons of adjustment error, pupil size, and microsaccade inhibition between different orientation distributions were analyzed using repeated measures ANOVA and paired-sample t-test. Multiple comparisons of post-hoc tests were corrected using Bonferroni correction. Multiple tests of beta values at multiple time points were corrected using FDR correction (Benjamini and Hochberg, 1995). Pearson correlation was calculated between adjustment error and pupil size across participants.

## Funding

This study was supported by National Science and Technology Innovation 2030 Major Program (2022ZD0204802 to F.F.) and the National Natural Science Foundation of China (32371086 to J.J., T2421004 and 31930053 to F.F.).

## Author contributions

J.J. and F.F. designed and supervised the work. J.J. and Z.K. acquired and analyzed the data and prepared the figures. All authors drafted and reviewed the manuscript.

## Competing interests

The authors declare that they have no competing interests.

## Data availability

Data supporting the main findings of the study are available at https://osf.io/92g7v/.

## Code availability

The code illustrating key analyses of the study can be found at https://github.com/perevo/pupilKZM.

## Notes

### Competing Interest Statement

The authors have declared no competing interest.

